# DeepImageTranslator V2: analysis of multimodal medical images using semantic segmentation maps generated through deep learning

**DOI:** 10.1101/2021.10.12.464160

**Authors:** En Zhou Ye, En Hui Ye, Maxime Bouthillier, Run Zhou Ye

## Abstract

**Introduction:** Analysis of multimodal medical images often requires the selection of one or many anatomical regions of interest (ROIs) for extraction of useful statistics. This task can prove laborious when a manual approach is used. We have previously developed a user-friendly software tool for image-to-image translation using deep learning. Therefore, we present herein an update to the DeepImageTranslator software with the addition of a tool for multimodal medical image segmentation analysis (hereby referred to as the MMMISA).

**Methods:** The MMMISA was implemented using the Tkinter library; backend computations were implemented using the Pydicom, Numpy, and OpenCV libraries. We tested our software using 4188 slices from whole-body axial 2-deoxy-2-[18F]-fluoroglucose-position emission tomography/computed tomography scans ([^18^F]-FDG-PET/CT) of 10 patients from the American College of Radiology Imaging Network-Head and Neck Squamous Cell Carcinoma (ACRIN-HNSCC) database. Using the deep learning software DeepImageTranslator, a model was trained with 36 randomly selected CT slices and manually labelled semantic segmentation maps. Utilizing the trained model, all the CT scans of the 10 HNSCC patients were segmented with high accuracy. Segmentation maps generated using the deep convolutional network were then used to measure organ specific [^18^F]-FDG uptake. We also compared measurements performed using the MMMISA and those made with manually selected ROIs.

**Results:** The MMMISA is a tool that allows user to select ROIs based on deep learning-generated segmentation maps and to compute accurate statistics for these ROIs based on coregistered multimodal images. We found that organ-specific [^18^F]-FDG uptake measured using multiple manually selected ROIs is concordant with whole-tissue measurements made with segmentation maps using the MMMISA tool.

## INTRODUCTION

Analysis of multimodal medical images (*e*.*g*., position emission tomography/magnetic resonance imaging [PET/MRI] and PET/computed tomography [PET/CT]) often requires the selection of one or many anatomical regions of interest (ROIs) for extraction of useful statistics [1-6]. The use of spherical or ellipsoid ROIs may be sufficient for large organs such as the liver and large muscle groups. However, for organs/tissues with complex shapes (*e*.*g*., the intestines and adipose tissues), manual ROI segmentation is not a scalable approach. One possible method is the use of deep learning for automated segmentation. Nevertheless, most deep learning pipelines for semantic image segmentation generate color-coded segmentation maps stored as image files, while most free software programs for medical image analysis (*e*.*g*., 3D-Slicer, OsiriX Lite, and AMIDE) cannot use these files to generate ROI statistics of multimodal images stored as DICOM files.

We have previously developed a user-friendly software tool for image-to-image translation using deep learning (DeepImageTranslator, described in [7], released at: https://sourceforge.net/projects/deepimagetranslator/). Therefore, we present herein an update to the DeepImageTranslator software with the addition of a tool for multimodal medical image segmentation analysis (hereby referred to as the MMMISA). We then demonstrate the use of the program for the measurement of 2-deoxy-2-[18F]fluoroglucose ([^18^F]-FDG) uptake by the lungs and subcutaneous adipose tissue using whole-body [^18^F]-FDG-PET/CT scans from the ACRIN-HNSCC-FDG-PET/CT database [8-10]. Furthermore, we also compare measurements performed using the MMMISA and those made with manually selected ROIs.

## METHODS

### Development of the MMMISA program

The MMMISA program presented herein was written in Python 3.8 and distributed under the GNU General Public License (version 3.0). The graphical user interface was developed using the Tkinter library, which is the most commonly used Python package for graphical user interface creation. Image analysis algorithms were implemented using the Pydicom, Numpy, and OpenCV libraries. The program is included as part of version 2 of the DeepImageTranslator software (https://sourceforge.net/projects/deepimagetranslator/) and is also available as a standalone program (https://sourceforge.net/projects/mmmisa/) for Windows. The source codes are available at: (https://github.com/runzhouye/MMMISA).

### PET/CT image dataset

Whole-body CT and FDG-PET images from 10 patients (numbers 001, 002, 003, 007, 008, 010, 012, 018, 019, and 027—which were the first 10 whole-body scans with the same windowing and devoid of significant radiographic artifacts) were downloaded from the ACRIN-HNSCC-FDG-PET/CT (ACRIN 6685) database [8, 9] via the Cancer Institute Archive [10]. We arbitrarily chose the ACRIN-HNSCC-FDG-PET/CT database since it was one of the few databases that contain coregistered whole-body PET and CT scans.

### Manual extraction of multimodal image data

CT and FDG-PET images were loaded into the AMIDE software [11]. For each patient, 11 spherical ROIs (10 mm diameter) in the subcutaneous adipose tissue and 3 spherical ROIs (50 mm diameter) in the lungs were drawn at different axial positions based on whole-body CT images. ROI statistics were subsequently generated for the coregistered PET images.

### Semantic image segmentation

Thirty-six axial slices were randomly chosen from the 4188 axial CT images from the 10 patients for manual semantic segmentation with the GIMP (GNU Image Manipulation Program) software of the background, lungs, bones, brain, subcutaneous and visceral adipose tissue, and other soft tissues by labelling these regions in black (RGB=[0,0,0]), yellow (RGB=[255,255,0]), white (RGB=[255,255,255]), cyan (RGB=[0,255,255]), red (RGB=[255,0,0]), green (RGB=[0,255,0]), and blue (RGB=[0,0,255]). Thirty-six training samples were considered more than sufficient since we have previously shown that models can be trained to achieve high accuracy with as little as 17 images [12]. CT image-segmentation map pairs were then loaded into the DeepImageTranslator software to train a deep convolutional neural network as previously described in Ye et al. [12] with 1000 training epochs. The final model was used to perform automatic semantic segmentation of the 4188 axial CT images from the 10 patients in less than 10 minutes. The generalizability of such an approach for automated segmentation has previously been shown to be excellent [12].

### Automated extraction of multimodal image data

For each patient, the original PET/CT scans were loaded into the MMMISA program along with the semantic segmentation maps produced by the convolutional neural network. In this study, we chose to extract FDG uptake from the lungs and subcutaneous adipose tissue by extracting regions of the model-generated segmentation maps containing yellow and red pixels, respectively using the MMMISA program. Lower and upper color threshold were set at (R,G,B) = (150,150,0) and (R,G,B) = (255,255,150), respectively, for the lungs, and (R,G,B) = (150,0,0) and (R,G,B) = (255,150,150), respectively, for the subcutaneous adipose tissue. ROI statistics were then generated for the FDG-PET scans using the MMMISA software.

### Statistical analyses

Statistical analyses were carried out using GraphPad Prism version 9. Pearson’s R values were computed for the correlation between organ-specific FDG uptake measured using multiple manually selected ROIs and FDG uptake determined using deep learning-generated segmentation maps.

### Data availability

The source code for the DeepImageTranslator is publicly available at: https://github.com/runzhouye/MMMISA

The compiled standalone software is available for Window10 at: https://sourceforge.net/projects/deepimagetranslator/ and https://sourceforge.net/projects/mmmisa/

The datasets generated during and/or analyzed during the current study are available at: https://doi.org/10.6084/m9.figshare.16800925

## RESULTS

### The MMMISA plugin for the DeepImageTranslator

The MMMISA program is included in version 2 of the DeepImageTranslator and is also available as a standalone software. The main window (**Fig.1**) allows for the user to visualize single- and dual-modality images written in the standard DICOM (Digital Imaging and Communications in Medicine) file format, the most commonly used file format in medical imaging. When images from a second modality are loaded into the program, they are automatically matched, along with the corresponding segmentation map, to the image of the first modality that is being currently displayed. When necessary, the program also performs image registration of modality 2 images based on modality 1 images through translation and/or stretching such that objects in both image sets overlap. This allows for simultaneous visualization of both image sets and segmentation maps.

**Fig.1:**
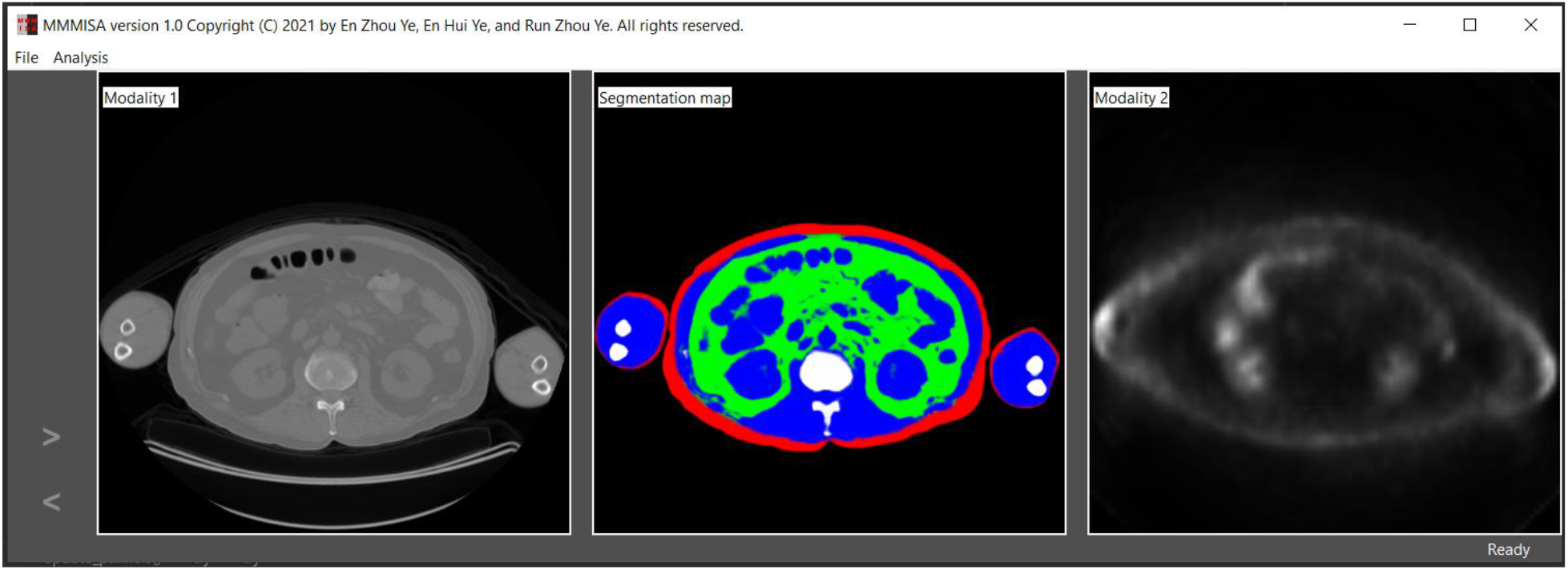
Main window of MMMISA, showing (from left to right), modality 1 (CT) images, segmentation maps generated with convolutional neural network, and modality 2 (PET) images.

A second, ROI selection, window (**Fig.2**) displays user settings for the extraction of ROIs based on pixel colors of the segmentation maps. Specific regions of the color-coded semantic segmentation maps can be extracted by setting lower and upper thresholds for the red, green, and blue color component values using the ROI selection window. The user can also choose to only include the left or right side of the patient for analysis, which can be useful in order to exclude the strong signals from of certain radiotracers injected into the left or right arm. When the “Apply” button is pressed, ROIs are generated based on the color thresholds using the segmentation maps and applied to corresponding slices of modality 1 and 2 images. The cropped images are then displayed in the main window for visualization.

**Fig.2:**
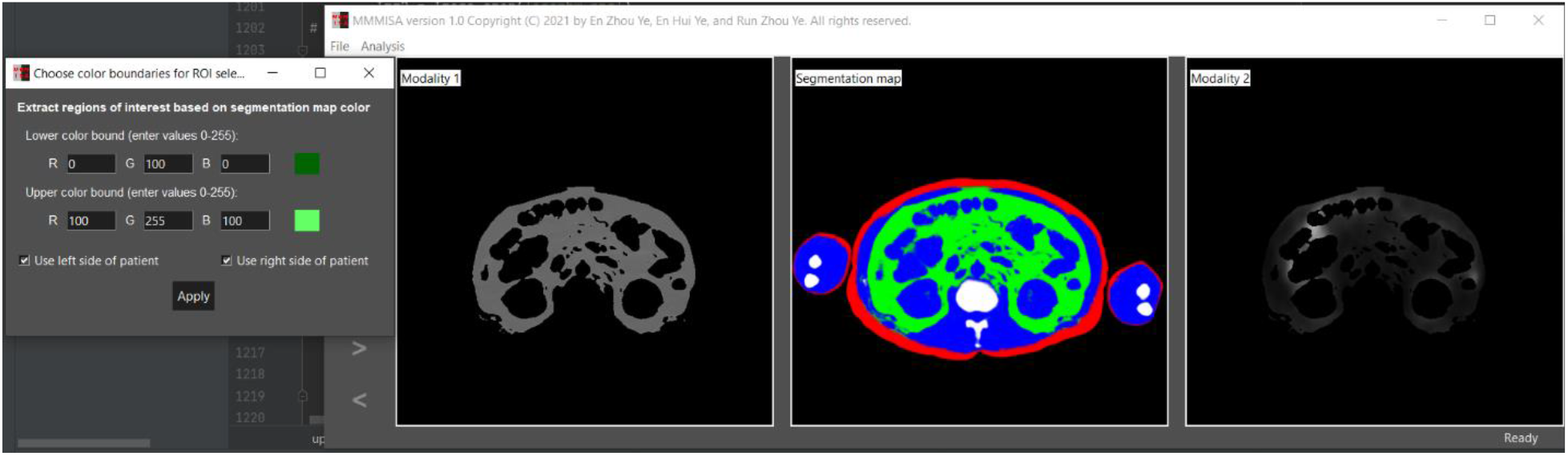
ROI selection window and main window with updated modality 1 and 2 images.

When “Save analysis” is selected, data will be extracted from modality 1 and 2 images, including the name of the scan, time at which each slice was produced, position of image slices, total area of the ROIs on each slice, total pixel values in the ROIs, average and standard deviation of values of pixels inside the ROIs, and pixel size. Results are then written in an excel file and stored under the user-designated directory.

### Semantic segmentation of PET/CT images

Segmentation results for images outside of the training set obtained with the convolutional neural network trained using the DeepImageTranslator were illustrated in **Fig.3**. Our final model was able to accurately segment the lungs, brain, bone, subcutaneous and visceral adipose tissue, and other soft tissues.

**Fig.3:**
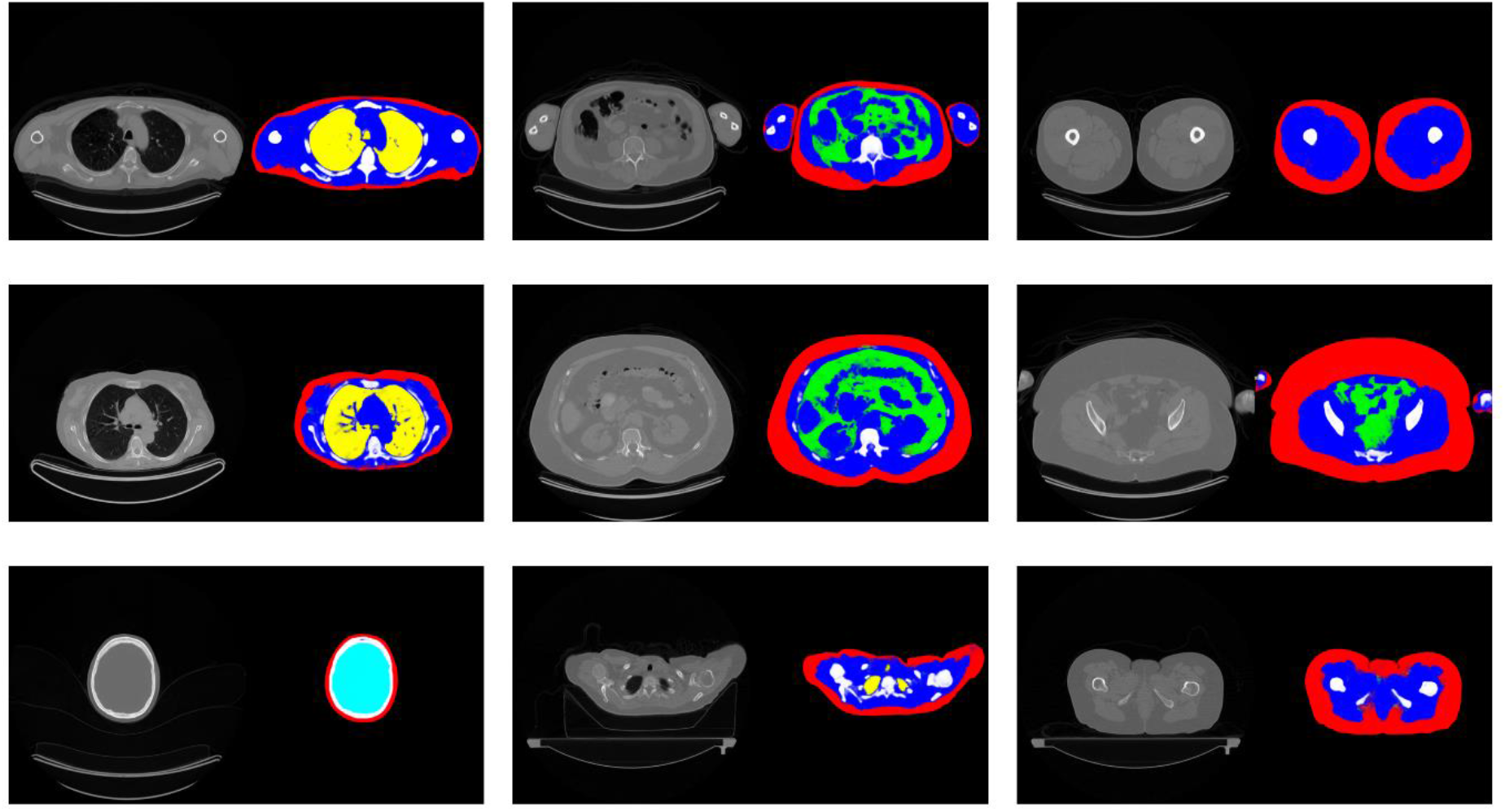
Pairs of CT images outside of the training set and sematic segmentation maps generated with deep convolutional neural network. The background, lungs, bones, brain, subcutaneous and visceral adipose tissue, and other soft tissues were labelled in black, yellow, white), cyan, red, green, and blue, respectively.

### Increase in number of manually selected ROIs increases accuracy of organ-specific FDG uptake approximations compared to true organ-specific FDG uptake measured using deep learning-generated segmentation maps

Next, we tested the concordance of organ-specific FDG uptake measured using multiple manually selected ROIs *versus* FDG uptake determined using deep learning-generated segmentation maps. In general, regardless the number of ROIs used, manually measured FDG uptake in the lungs and subcutaneous adipose tissue was well correlated with that calculated with segmentation maps using the MMMISA program (**Fig.4**). For subcutaneous adipose tissue FDG uptake, the correlation coefficient and the -log of the P-value increased sharply once values from more than 4 ROIs were combined (**Fig.4A**). Increase in measurement accuracy (determined by the correlation coefficient) through increasing numbers of manually selected ROIs plateaued after more than 8 ROIs were used. Nevertheless, the P-value of the correlation between manual measurement and that using segmentation maps continued to decrease when more ROIs were used (**Fig.4B**). Similar results were obtained for the measurement of FDG uptake in the lungs (**Fig.4C-D**).

**Fig.4:**
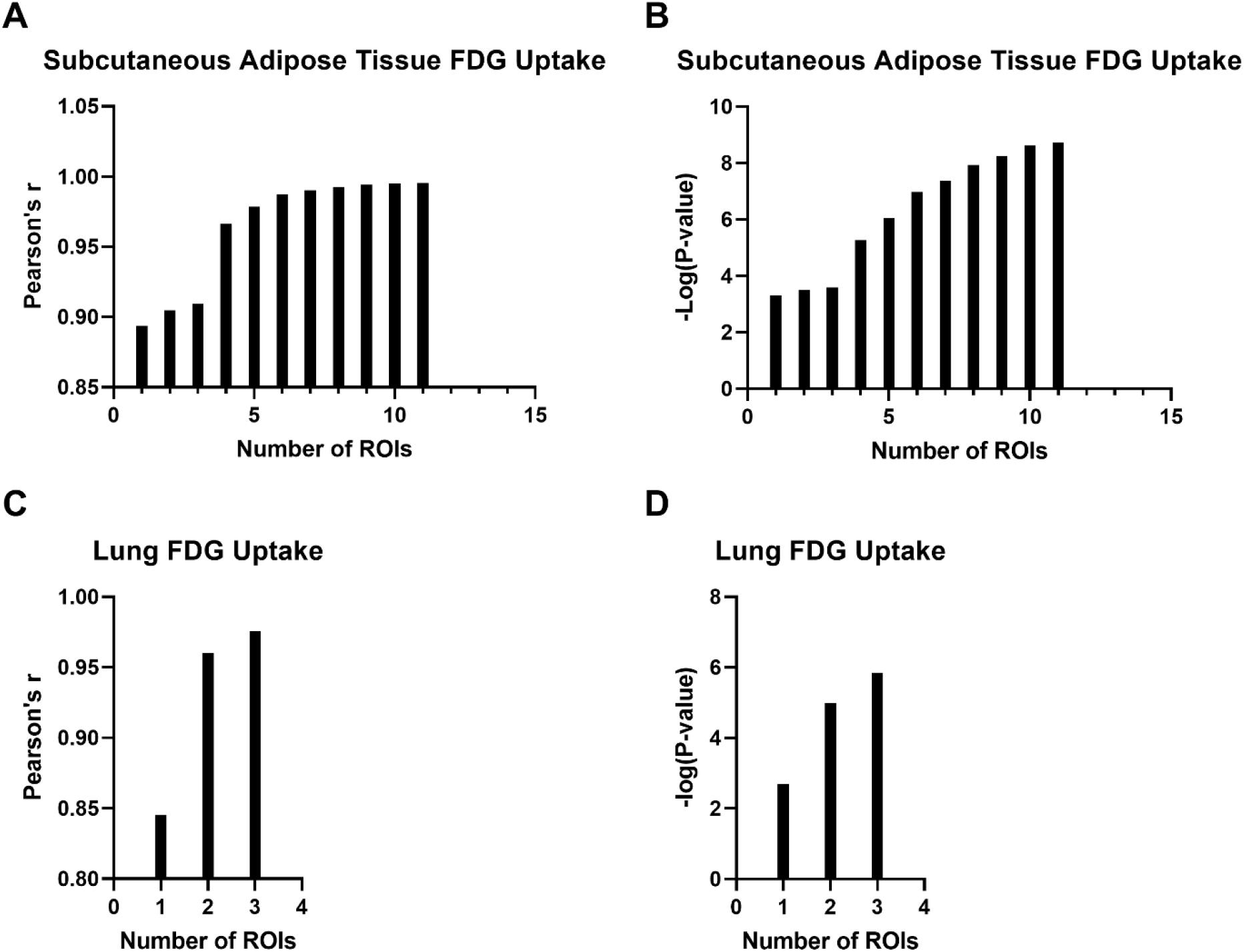
Correlation coefficient (**A** and **C**) and P-value (**B** and **D**) for the association between organ-specific FDG uptake measured using multiple manually selected ROIs and FDG uptake determined using deep learning-generated segmentation maps, as a function of number of manually selected ROIs, for the subcutaneous adipose tissue (**A** and **B**) and lungs (**C** and **D**). ROI: region of interest.

## DISCUSSION

In recent years, numerous open-source software tools have been reported in the field of medical image processing [13-17]. One growing area of development is the popularization of deep learning methods through the creation of user-friendly tools with a graphical interface. Nevertheless, most deep learning pipelines for semantic image segmentation generate color-coded segmentation maps stored as image files, while most free software programs for medical image analysis cannot use these files to generate ROI statistics of multimodal images stored as DICOM files.

Nonetheless, selection of ROIs is an important aspect of *in vivo* metabolic studies involving PET/CT imaging [18-21]. In particular, measurements of volume and radiotracer uptake of adipose tissues of different regions may prove to be important for future studies on the metabolic syndrome, as hypertrophic obesity is related to changes in adipose tissue distribution and alterations in metabolic endpoints [22, 23].

Therefore, we have presented herein an update to the DeepImageTranslator software [7] by including a tool for multimodal medical image segmentation analysis based on semantic segmentation maps generated using a deep convolutional neural network. Our program can be accessed through a graphical interface and allows users to extract ROI statistics of multimodal images (*e*.*g*., PET/CT and PET/MRI) based on color-coded semantic segmentation maps. We showed that organ-specific FDG uptake measured using multiple manually selected small, spherical ROIs is concordant with whole-tissue measurements made with segmentation maps using the MMMISA program. Furthermore, we found that increase in number of manually selected ROIs increases the accuracy of organ specific FDG uptake approximations. Therefore, our pipeline constitutes a simple, automated, and scalable approach to obtain ROI statistics using multimodal scans. Although the accuracy of the neural network would never surpass the accuracy of manually labelled segmentation maps used for model training, our approach would eventually greatly simplify the task of researchers and radiologists performing whole-body semantic segmentation of multimodal tomography data.

## DISCLOSURES

The authors declare no competing interests.

## AUTHOR CONTRIBUTIONS

Software development: EZY, EHY, and RZY. Statistical analyses: EHY. Interpretation and manuscript drafting: EZY, MB, and RZY.

## Notes

https://doi.org/10.6084/m9.figshare.16800925

https://sourceforge.net/projects/mmmisa/

https://github.com/runzhouye/MMMISA

### Competing Interest Statement

The authors have declared no competing interest.

